# *Drosophila* insulin-like peptide *dilp1* increases lifespan and glucagon-like *Akh* expression epistatic to *dilp2*

**DOI:** 10.1101/380410

**Authors:** Stephanie Post, Sifang Liao, Rochele Yamamoto, Jan A. Veenstra, Dick R. Nässel, Marc Tatar

**Affiliations:** Department of Molecular Biology, Cell Biology and Biochemistry, Providence, RI, Brown University, United States of America.; Department of Ecology and Evolutionary Biology, Providence, RI, Brown University, United States of America.; Department of Zoology, Stockholm University, S-10691 Stockholm, Sweden.; Institut de Neurosciences Cognitives et Intégratives d’Aquitaine (CNRS UMR5287), University of Bordeaux, Pessac, France.

**Keywords:** Drosophila, insulin-like peptide, dilp1, dilp2, insulin/IGF signaling, Akh, aging

## Abstract

Insulin/IGF signaling (IIS) regulates essential processes including development, metabolism, and aging. The *Drosophila* genome encodes eight insulin/IGF-like peptide *(dilp)* paralogs, including tandem-encoded *dilp1* and *dilp2.* Many reports show that longevity is increased by manipulations that decrease DILP2 in adults. In contrast, *dilp1* is expressed primarily in pupal stages, but also during adult reproductive diapause, although we find that *dilp1* is also highly expressed in adult *dilp2* mutants under non-diapause conditions. The inverse expression of *dilp1* and *dilp2* suggests these genes interact to regulate aging. Here, we study *dilp1* and *dilp2* single and double mutants to describe epistatic and synergistic interactions affecting longevity, metabolism and adipokinetic hormone (AKH), a functional homolog of glucagon. Mutants of *dilp2* extend lifespan and increase *Akh* mRNA and protein in a *dilp1-*dependent manner. Loss of *dilp1* alone has no impact on these traits, whereas transgene expression of *dilp1* increases lifespan in *dilp1-dilp2* double mutants. On the other hand, *dilp1* and *dilp2* redundantly interact to control circulating sugar, starvation resistance and compensatory *dilp5* expression. These later interactions do not correlate with patterns for how *dilp1* and *dilp2* affect longevity and AKH. Thus, repression or loss of *dilp2* slows aging because its depletion induces *dilp1,* which acts as a pro-longevity factor. Likewise, *dilp2* regulates *Akh* through epistatic interaction with *dilp1. Akh* and glycogen affect aging in *C. elegans* and *Drosophila,* suggesting that *dilp2* modulates lifespan via *dilp1* and in part by regulating *Akh.* Whether DILP1 acts as an insulin receptor agonist or inhibitor remains to be resolved.

## Introduction

Insulin/IGF signaling (IIS) is a fundamental pathway that regulates aging, development, metabolism, growth and other critical systems. The *Drosophila melanogaster* genome encodes several insulin-like peptide genes (*dilps*) that signal through a single insulin-like receptor *(InR)* and one apparent set of signal transduction elements (Brogiolo et al., 2001; Colombani, Andersen, & Leopold, 2012; Garofalo, 2002; Grönke, Clarke, Broughton, Andrews, & Partridge, 2010). Among their physiological functions, *dilps* regulate aging: mutation of *dilp2* alone is sufficient to extend lifespan, although longevity is somewhat further extended in a *dilp2-3,5* triple mutant (Grönke et al., 2010). How specific *dilps* modulate aging is not understood. Here, we demonstrate that *dilp1* is upregulated in the absence of *dilp2,* that *dilp1* expression is required for loss of *dilp2* to slow aging, and that exogenous expression of *dilp1* in a *dilp1-2* mutant is sufficient to extend lifespan.

The *dilp1* gene is encoded approximately 1.2kb upstream of *dilp2,* potentially as a result of a tandem duplication event (Tatar, Bartke, & Antebi, 2003). The two paralogs are expressed in different developmental and life history stages. *Dilp2* is initially expressed in embryos and then throughout larval instar stages (Brogiolo et al., 2001; Slaidina, Delanoue, Grönke, Partridge, & Leopold, 2009). Pupae show decreased expression of *dilp2,* but the ligand is again highly expressed in adults. In contrast, during normal development, *dilp1* is only expressed at high levels in the pupal stage, when *dilp2* mRNA is minimal (Slaidina et al., 2009). While their timing is distinct, *dilp1* and *dilp2* are both expressed in median neurosecretory cells of the *Drosophila* brain, the insulin-producing cells (IPCs) analogous to mammalian pancreatic β-cells (Brogiolo et al., 2001; Broughton et al., 2005; Y. Liu, Liao, Veenstra, & Nässel, 2016; Rulifson, Kim, & R., 2002).

The function of *dilps* in aging has been best studied for *dilp2, dilp3, dilp5* and *dilp6* (Bai, Kang, & Tatar, 2012; Broughton et al., 2010; Grönke et al., 2010). Mutation of *dilp2* alone, or *dilp2, dilp3* and *dilp5* together, extend lifespan: the normal function of these ligands appears to promote processes permissive to aging (Grönke et al., 2010). On the other hand, induction of *dilp6* in fat body promotes longevity, perhaps because this decreases DILP2 secreted from the IPCs (Bai et al., 2012). Similarly, increased *FOXO* expression in head fat body and increased JNK activity in IPCs extends lifespan, perhaps again because these manipulations decrease *dilp2* expression in the IPCs (Hwangbo, Gershman, Tu, Palmer, & Tatar, 2004; Wang, Bohmann, & Jasper, 2005). Across these studies, there has been no attention to *dilp1,* which appears to only be produced in the adult IPCs during the state of adult reproductive diapause (Liu et al., 2016). *Drosophila* diapause, however, is strongly associated with slow or negligible aging (Tatar & Yin, 2001). The positive association between *dilp1* and diapause survival suggests that this enigmatic insulin hormone may possess unusual functions in the control of aging.

Understanding how these various *dilps* regulate aging is complicated by the fact that genetic or RNAi reduction of any one *dilp* gene induces compensatory expression in other *dilp* genes. For instance, a *dilp2* mutant increases expression of *dilp3* and *dilp5* and these ligands consequently stimulate insulin/IGF signaling (Grönke et al., 2010). Such compensation makes it difficult to dissect the role of signaling downstream of the fly insulin/IGF receptor in aging. Complex compensation and interaction is also known for *C. elegans* insulin-like gene paralogs (Fernandes de Abreu et al., 2014). For example, *ins-6* is upregulated in an *ins-23* mutant, and these two paralogs interact epistatically to regulate lifespan (Fernandes de Abreu et al., 2014). Notably, *C. elegans ins-18* and *ins-23* are proposed to act as insulin-like receptor antagonists that additively regulate dauer formation and favor longevity (Matsunaga, Matsukawa, Iwasaki, Nagata, & Kawano, 2018). Some interactions are established for *Drosophila* insulin paralogs, such as the inverse regulation of aging by *dilp6* and *dilp2* (Bai et al., 2012), but the functional relationships between *dilp2* and other *dilps* have not been characterized.

The relationship between *dilp1* and *dilp2* may represent in *Drosophila* a longevity regulatory system similar to that suggested with *C. elegans* paralogs *ins-6* and *ins-23* (Fernandes de Abreu et al., 2014). We find that *dilp1* is strongly upregulated in *dilp2* mutants, consistent with *dilp1* serving a role in diapause conditions where it might regulate metabolism and slow aging. To test this model, we generated *dilp1-2* double mutants to complement similarly constructed *dilp1* and *dilp2* single mutants (Gronke et al., 2010). As previously reported, *dilp2* mutants are long-lived. We now see that *dilp1* mutants have wildtype longevity as do *dilp1-dilp2* double mutants; thus loss of *dilp1* fully rescues the extended longevity of *dilp2* to wildtype. We find that *dilp1* is also downstream of *dilp2* in the control of *Drosophila* adipokinetic hormone (AKH), the functional homolog of mammalian glucagon. We confirmed the positive role of *dilp1* upon longevity and AKH by transgene *dilp1* expression in a *dilp1-dilp2* double mutant. In contrast to longevity and AKH, *dilp1* and *dilp2* do not control other physiological traits (e.g. hemolymph sugar, starvation resistance and glycogen) in this epistatic manner, suggesting that these phenotypes are not regulated through the same mechanisms by which these insulin-like peptides modulate aging. Our data together reveal a novel pathway by which a unique insulin-like ligand, DILP1, positively regulates longevity.

## Results

Studies on the control of aging by IIS in *Drosophila* have measured *dilp2, dilp3* and *dilp5* mRNA or protein (Alic, Hoddinott, Vinti, & Partridge, 2011; Broughton et al., 2010; Hwangbo et al., 2004). While *dilp1* of the adult IPC is not observed in non-diapause conditions, we sought to characterize its expression in *dilp* mutants known to extend lifespan. In wildtype adult females, *dilp1* mRNA is considerably lower than that of *dilp2* (Fig 1A). Strikingly, *dilp1* mRNA is elevated about 14-fold in *dilp2* mutants relative to its expression in wildtype (Fig 1B), while there is little compensatory expression of *dilp2* in *dilp1* mutants (Fig 1C). *Dilp2* appears to repress *dilp1,* and we propose that *dilp1* may function in the absence of *dilp2* to regulate metabolism and aging.

**Figure 1.**
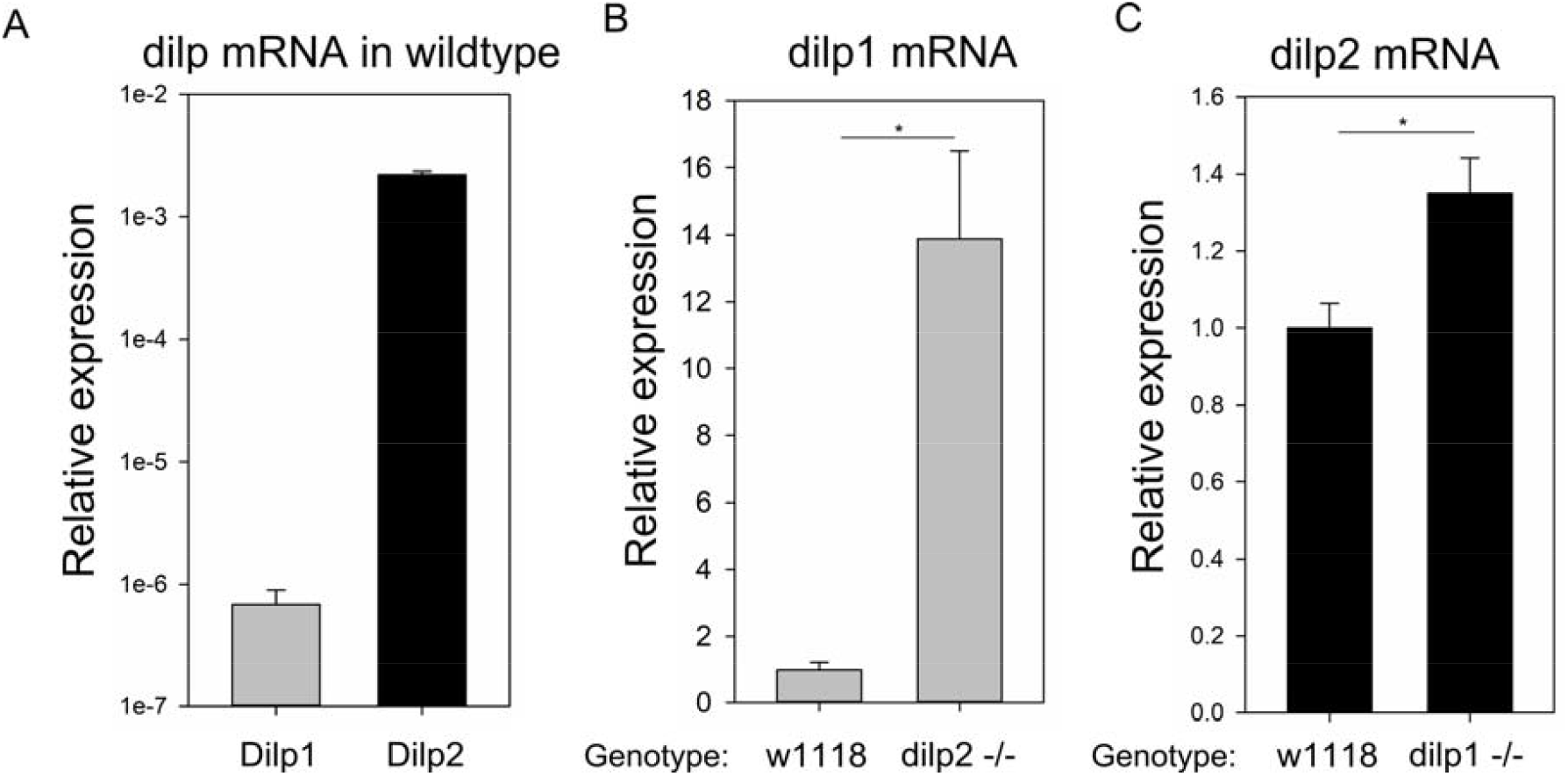
*Dilp1* mRNA is induced by depletion of *dilp2.* RNA from 7-10 day old female adult flies was assayed by q-RT-PCR. n=6 per genotype. (A) *Dilp1* expression is 100-fold lower than *dilp2* expression in wildtype flies. (B) *Dilp1* expression increases 100-fold in *dilp2* mutant flies compared to wildtype flies, t test p<0.001. (C) *Dilp2* expression increases 2-fold in *dilp1* mutant flies compared to wildtype flies, t test p=0.005.

### Epistasis analysis of lifespan

Adult *dilp2* mutants have elevated blood sugar and extended lifespan (Grönke et al., 2010). To test if these phenotypes require the expression of *dilp1,* we generated a *dilp1-dilp2* null double mutant by homologous recombination, in parallel with matching *dilp1* and *dilp2* null single mutant knock-outs (Fig. S1, Supporting Information). If the functions of *dilp1* are redundant to those of *dilp2,* we would expect the *dilp1-dilp2* double mutants to have greater longevity and higher blood sugar than either single mutant. Alternatively, if the functions of *dilp1* are downstream of *dilp2,* we expect the *dilp1-dilp2* double mutants to have wildtype lifespan and metabolism.

Our new null allele of *dilp2* increases lifespan by about 20-30% (Fig 2A, Fig S2D, Supporting Information), confirming previous observations (Grönke et al., 2010). Null mutation of *dilp1* has no effect on adult survival, again as previously reported (Grönke et al., 2010). Remarkably, survival of the *dilp1-dilp2* double null mutant is indistinguishable from wildtype or the *dilp1* mutant (Fig 2A-B), revealing a classic epistatic interaction between *dilp1* and *dilp2* in the control of longevity.

**Figure 2.**
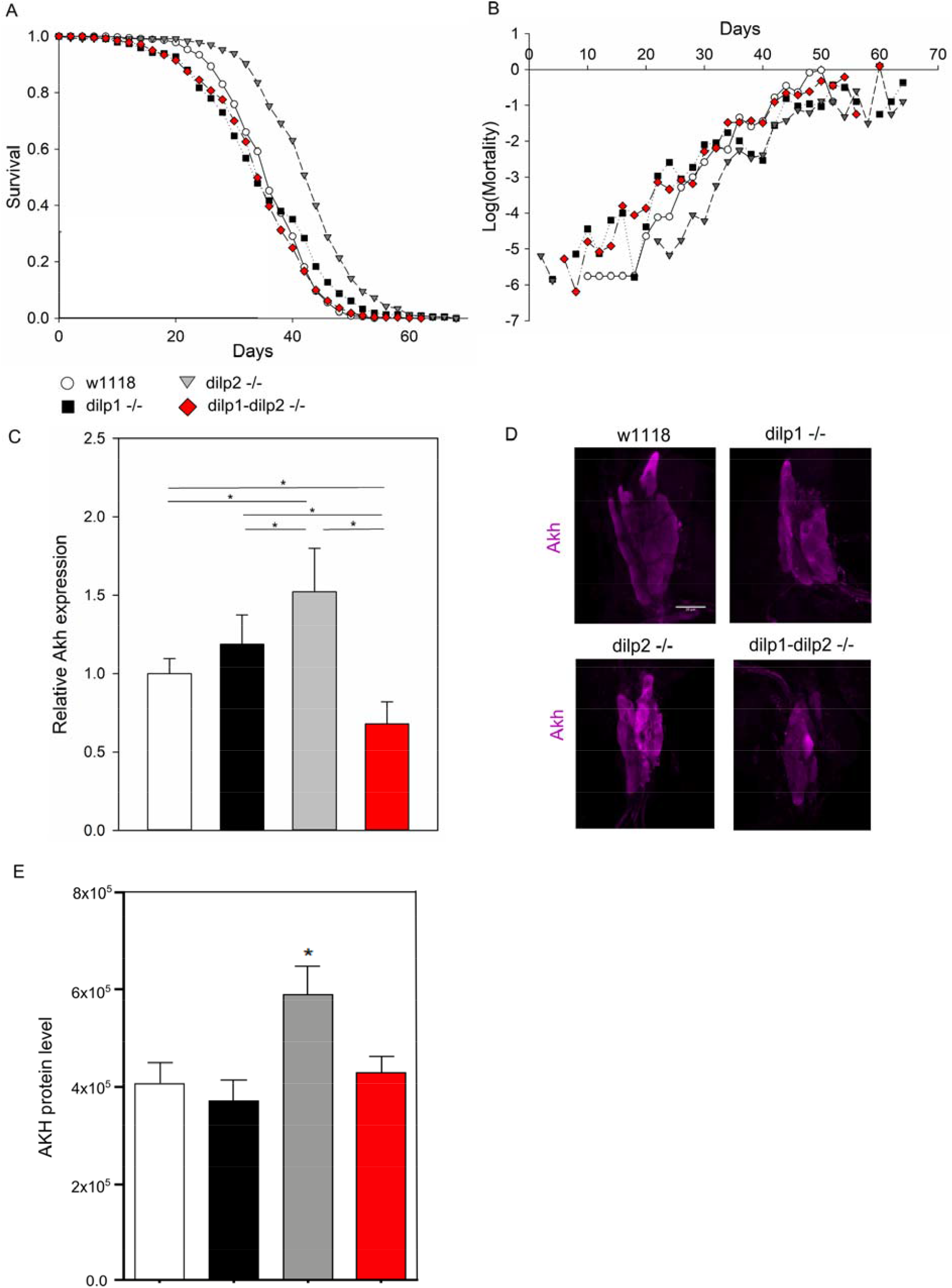
A *dilp1* mutation suppresses aging and *Akh* phenotypes of mutant *dilp2.* (A) *Dilp2* mutants but not double mutants are long-lived, Cox hazard analysis p<0.0001, chi^2^=201, n=341-365 per genotype. (B) *Dilp2* mutants but not double mutants have decreased mortality. (C) *Dilp2* mutants but not double mutants have increased *Akh* expression. RNA from 7-10 day old female adult flies was assayed by q-RT-PCR, n=9 per genotype. Two-way ANOVA dilp1 p=<0.001, dilp2 p=0.921, dilp1 x dilp2 p<0.001. (D) *Dilp2* mutants but not *dilp1* or double mutants have increased AKH immunolabeling in corpora cardiaca from 6-7 day old female flies. Representative images shown. (E) Quantification of AKH immunolabeling, n=9-14 samples from 3 replicates, ANOVA * p<0.05.

### Epistasis analysis of adipokinetic hormone

*Dilp1* is only normally expressed in adults during reproductive diapause, a slow-aging stage associated with many metabolic changes, including activity of adipokinetic hormone (AKH), the functional homolog of mammalian glucagon (Kubrak, Kucerova, Theopold, & Nässel, 2014; Kucerova et al., 2016; Y. Liu et al., 2016). Accordingly, we studied how *dilp1* and *dilp2* affect AKH through genetic analysis of single and double mutants. *Akh* mRNA is increased in *dilp2* mutants, is similar to wildtype in *dilp1* mutants, and is restored to wildtype levels in the *dilp1-dilp2* double mutants (Fig 2C). Physiologically, insulin regulates glucagon at the level of mRNA and protein (Pearson, Unger, & Holland, 2016). We therefore examined AKH immunostaining in the adult corpora cardiaca (CC). AKH peptide in the CC is increased in *dilp2* mutants, and is similar to wildtype in *dilp1* and in *dilp1-dilp2* double mutants (Fig 2D-E). These data suggest that *dilp1* is epistatically downstream of *dilp2* in pathways to regulate lifespan and AKH – *dilp1* expression is required for *dilp2* to modulate these phenotypes.

### Epistasis analysis of developmental and metabolic traits

*Drosophila* insulin, including *dilp1* and *dilp2,* affect many additional traits, including body weight and metabolism (Grönke et al., 2010). Similar to the epistatic interactions observed for lifespan and AKH, body mass was decreased in *dilp2* mutants (as previously reported (Grönke et al., 2010)), but similar to wildtype in *dilp1* mutants and in *dilp1-dilp2* double mutants (Fig 3A). In contrast, hemolymph (blood) sugar concentrations in single mutants of *dilp1* and *dilp2* are similar to those seen in wildtype, while the *dilp1-dilp2* double mutant has elevated hemolymph sugar (Fig 3B): for this trait, these insulin paralogs appear to have parallel, redundant functions (Table 1). On the other hand, glycogen content is equally decreased by both single mutants and the double mutant, indicating that both *dilp1* and *dilp2* are required to maintain the titer of this energy storage molecule (Fig 3C).

**Table 1.**
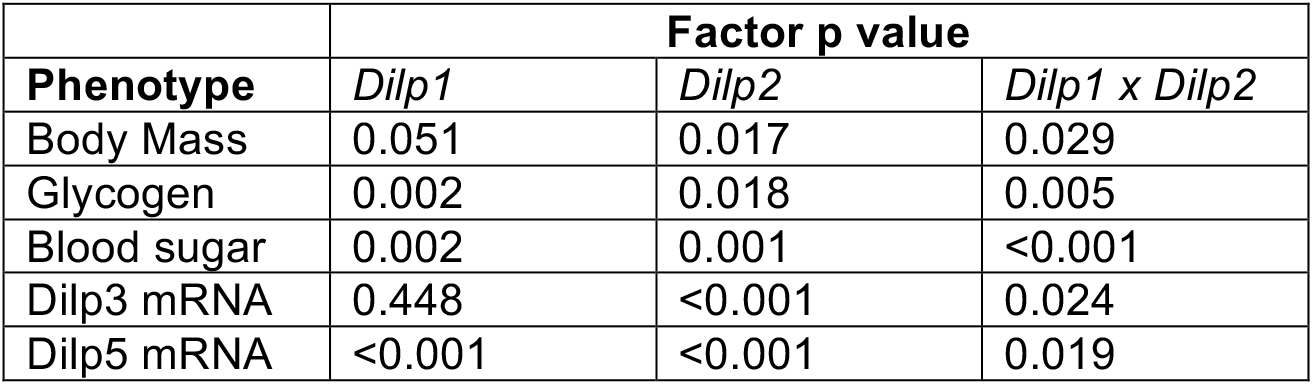
Two-way ANOVA p values for dilp1-dilp2 epistasis between single and double mutants

**Figure 3.**
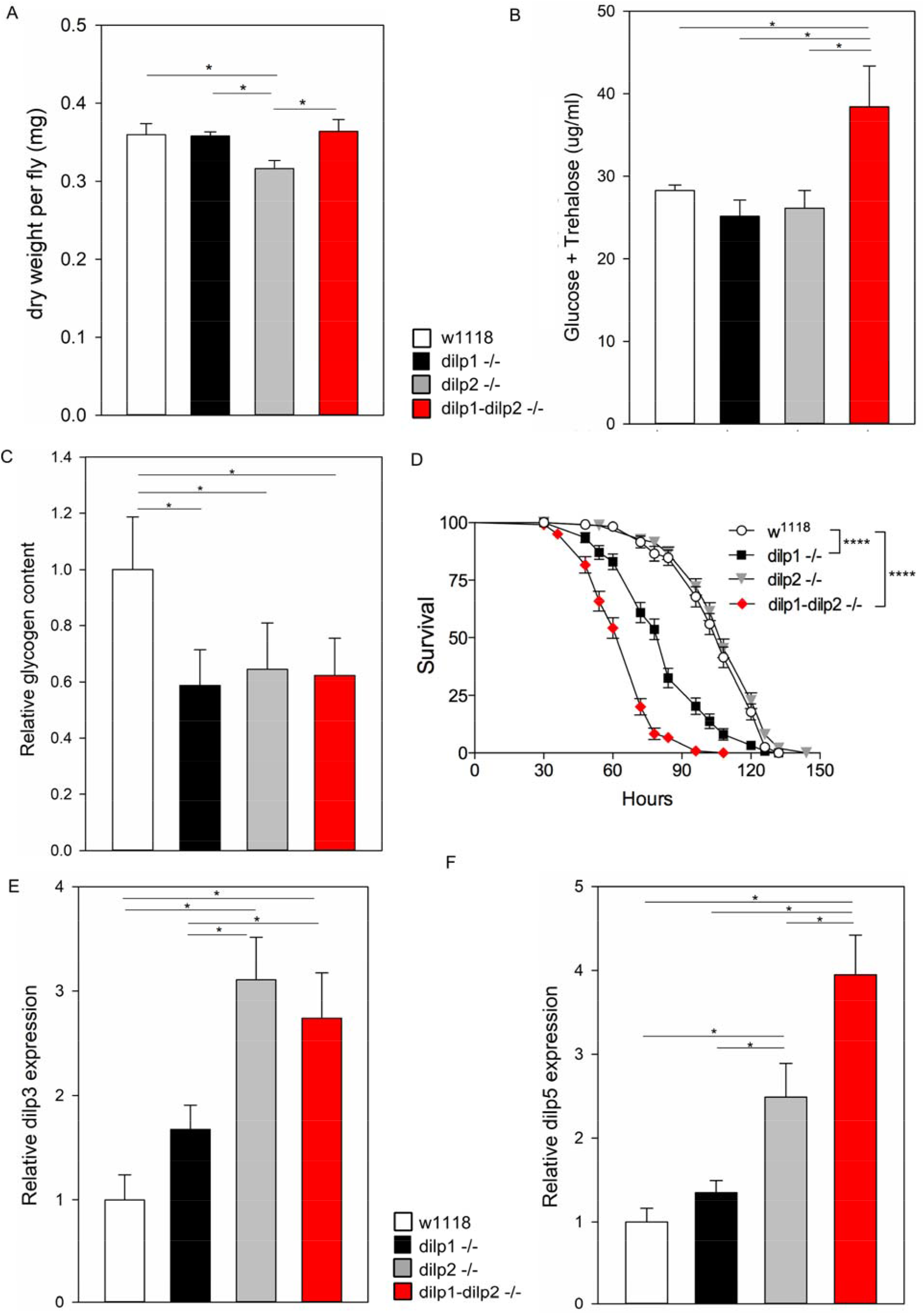
*Dilp1* and *dilp2* interact to regulate metabolism, physiology and dilp compensation. Female flies 7-10 days old were assayed. For two-way ANOVA statistics, refer to Table 1. (A) Body size is decreased in *dilp2* mutants, n=5 per genotype, 22-44 flies per replicate. Hemolymph sugar levels (B) and glycogen levels (C) differ significantly between single and double mutants, n=5 per genotype. (D) Starvation sensitivity differs significantly between single and double mutants, n=118-150 flies from 3 replicates, Log-rank (Mantel-Cox) **** p<0.0001. *Dilp3* (E) and *dilp5* (F) expression compensate for *dilp1* and *dilp2* differently, n=3-6 per genotype.

Many longevity-extending IIS manipulations increase resistance to starvation (Clancy et al., 2001; Grönke et al., 2010). Unexpectedly, our long-lived *dilp2* null genotype has starvation survival similar to wildtype, while *dilp1* mutants and *dilp1-dilp2* double mutants are more sensitive to starvation (Fig 3D). These data suggest that *dilp1,* which is increased during nonfeeding developmental stages (Y. Liu et al., 2016), may be required for starvation survival by inducing catabolism of nutrients such as glycogen, perhaps through the action of AKH. *Dilp3* mRNA is increased in *dilp2* mutants and similarly in *dilp1-dilp2* double mutants, but not significantly increased in *dilp1* mutants (Fig 3E). *Dilp5* mRNA is increased to a greater extent in *dilp1-dilp2* double mutants relative to its increase in either single mutant, representing synergistic genetic interaction between *dilp1* and *dilp2* (Fig 3F). We observed no induction or repression of mRNA for *dilp6, dilp7* and *dilp8* in single and double mutants of *dilp1* and *dilp2* (Fig S2A-C, Supporting Information). Fecundity is not significantly different among wildtype, single and double *dilp1* and *dilp2* mutants for adult females one or three weeks old, although at two weeks old, *dilp2* mutants lay slightly more eggs per day than the other genotypes (Fig S2E, Supporting Information). Finally, survival measured from egg through pupal stage was 50% less in *dilp1-dilp2* double mutants relative to wildtype and to single mutants, suggesting that these insulin loci have redundant functions in egg-to-pupal viability (Fig S2F, Supporting Information). In sum, lifespan, body size and AKH are mediated by classic genetic epistasis between *dilp1* and *dilp2,* where *dilp1* functions downstream of dilp2. Other measured phenotypes are jointly regulated by *dilp1* and *dilp2,* in some cases by redundant functions of the ligands and in other through synergistic functional interactions.

### Epistatic analysis of insulin/IGF and juvenile hormone signaling

To understand how *dilp1* is required to extend longevity, we evaluated insulin/IGF signal (IIS) transduction and juvenile hormone (JH) signaling in single and double *dilp1* and *dilp2* mutants. Insulin ligands in *Drosophila* are described to induce phosphorylation of Akt and ERK, which in turn regulate activity of transcription factors including FOXO. Here, we measured Akt and ERK phosphorylation from thorax tissue, which primarily consists of flight muscle (Fig 4A-C). While loss of *dilp1* had no impact on Akt phosphorylation, loss of *dilp2* *increased* Akt phosphorylation in single and double mutants, suggesting that compensatory expression of other *dilps (dilp3* and *dilp5)* are sufficient to maintain this branch of IIS in the absence of *dilp2.* In contrast, ERK phosphorylation in thorax is reduced in *dilp2* mutants, is unaffected in *dilp1* mutants, and is restored to wildtype levels in the *dilp1-dilp2* double mutant. *Dilp1* and *dilp2* interact epistatically to control ERK phosphorylation. This pattern correlates with the epistatic interaction we observe for *dilp1* and *dilp2* in the control of longevity, and we note that ERK has been implicated in how IIS controls aging downstream of the insulin receptor substrate *chico* (Slack et al., 2015).

**Figure 4.**
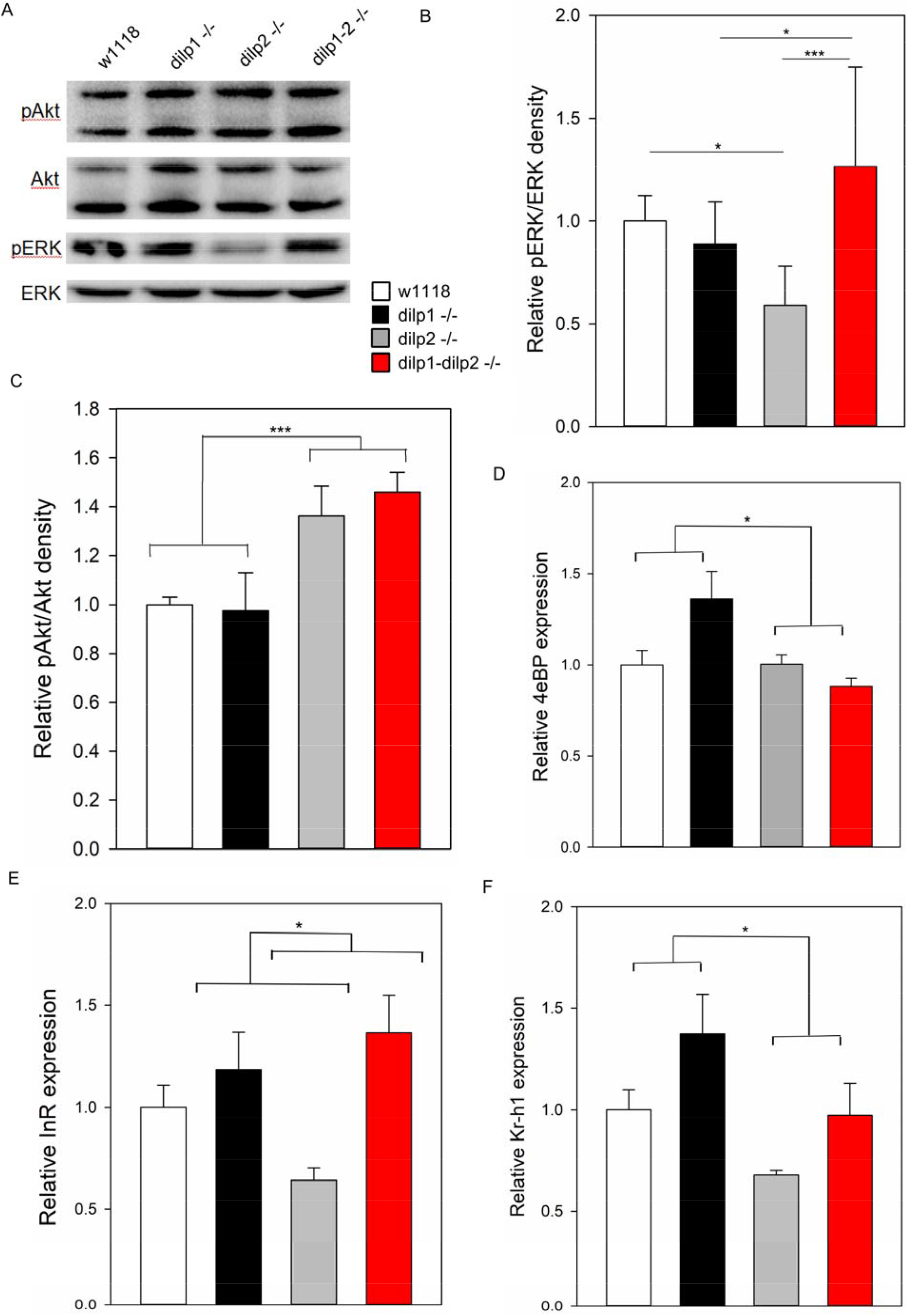
Components of insulin/IGF and JH signaling regulated by *dilp1* and *dilp2.* (A) Dissected thorax pAkt is increased in *dilp2* and double mutants, and pERK is decreased in *dilp2* mutants in a *dilp1*-dependent manner, representative blot shown. (B) Quantification of thorax pERK/ERK phosphowesterns, two-way ANOVA dilp1 x dilp2 p=0.003, n=6 per genotype. (C) Quantification of thorax pAkt/Akt phosphowesterns, two-way ANOVA dilp2 factor p<0.001, dilp1 x dilp2 N.S, n=6 per genotype. (D) *4eBP* expression is not elevated in *dilp2* mutants, two-way ANOVA dilp1 x dilp2 p=0.01, n=7-9 per genotype. (E) *InR* expression is not elevated in *dilp2* mutants, but differentially expressed between single and double mutants, two-way ANOVA dilp1 x dilp2 p=0.054, n=7-9 per genotype. (F) *Kr-h1* expression is decreased in *dilp2* mutants, two-way ANOVA dilp1 p=0.02, dilp2 p=0.013, dilp1 x dilp2 N.S., n=8-9 per genotype.

Reduced IIS extends *Drosophila* lifespan in part through activating the FOXO transcription factor (Bai, Kang, Hernandez, & Tatar, 2013; Hwangbo et al., 2004; Min, Yamamoto, Buch, Pankratz, & Tatar, 2008). Accordingly, we measured FOXO activation in *dilp1* and *dilp2* single and double mutants by measuring expression of FOXO transcriptional targets *4eBP* and *InR* from whole animals (Fig 4D-E). Neither FOXO target was elevated in the long-lived *dilp2* single mutant. Thus, with our current mutants we find no association between longevity and elevated *4eBP* or *InR* expression, suggesting that FOXO activation is not responsible for how reduced *dilp2* slows aging.

Juvenile hormone (JH) is an insect terpenoid hormone produced by the corpora allata and documented to modulate how IIS impacts aging, where eliminating adult JH production is sufficient to extend lifespan (Tatar et al., 2001; Yamamoto, Bai, Dolezal, Amdam, & Tatar, 2013). JH controls transcriptional programs by regulating expression of the transcription factor Kruppel-homolog 1 (Kr-h1) (S. Liu et al., 2018; Minakuchi, Zhou, & Riddiford, 2008). Unlike FOXO, the expression of *Kr-h1* mRNA (and thus JH activity) follows a pattern consistent with the epistatic interaction between *dilp1* and *dilp2* as seen for longevity: *Kr-h1* is reduced in *dilp2* mutant and restored to wildtype levels in the *dilp1-dilp2* double mutant, while *dilp1* single mutants tend to have greater *Kr-h1* mRNA than wildtype (Fig 4F). *Dilp1* thus appears to normally repress JH activity, and we conjecture that this may be sufficient for *dilp1* to extend longevity when induced in *dilp2* mutants.

### Overexpression of dilp1 to rescue phenotypes in dilp1-dilp2 double mutant

We sought to confirm our key inferences from the epistasis analyses by rescuing phenotypes in the double mutant by exogenously expressed *dilp1.* We generated and validated a new UAS-*dilp1* stock to express this insulin protein using GAL4-drivers (Fig S4A, Supporting Information). To test whether *dilp1* is sufficient to slow aging and increase *Akh* expression, we induced UAS-*dilp1* in the line containing *dilp1-dilp2* null mutations. Driving UAS-*dilp1* in IPCs with *dilp2-GAL4* (Fig 5A-B) or in all neurons with the RU486 inducible GeneSwitch elav-GSGal4 (Fig 5C-D) significantly extended lifespan by consistently decreasing age-specific mortality. Likewise, *Akh* mRNA was elevated by *dilp1* transgene expression when driven by *dilp2-GAL4* in the double mutant background (Fig 5E).

**Figure 5.**
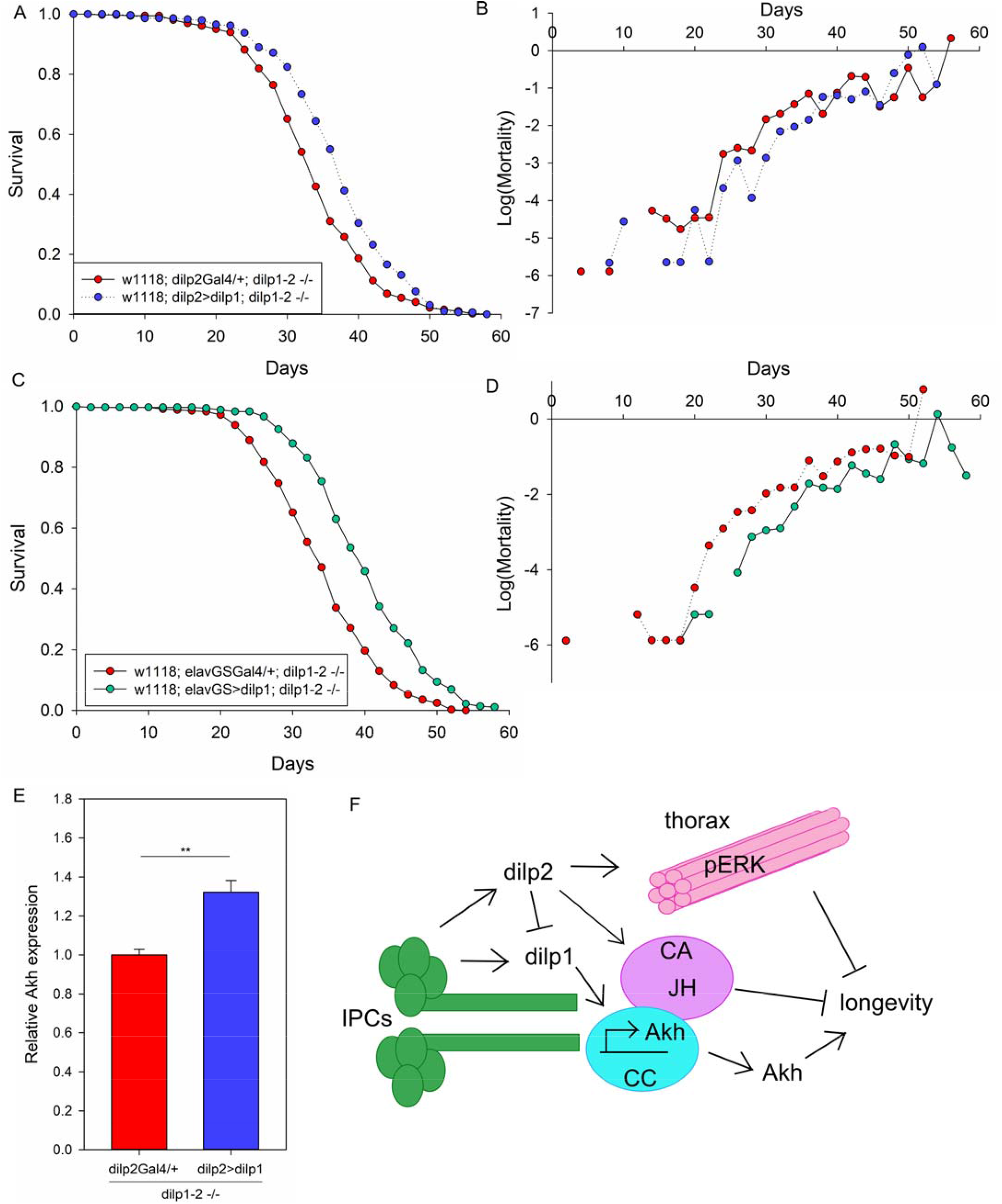
*Dilp1* expression in double mutants rescues longevity and AKH. (A) Lifespan is extended by *dilp2>dilp1* overexpression rescue in the *dilp1-dilp2* double mutant background compared to dilp2-GAL4/+ controls, Cox hazard analysis p<0.0001, chi^2^=25.8, n=289-364 per genotype. (B) Mortality is decreased in *dilp2>dilp1* overexpression rescue in the *dilp1-dilp2* double mutants compared to *dilp2-GAL4/+* controls. (C) Lifespan is extended by *elav-GS>dilp1* overexpression in the *dilp1-dilp2* double mutant background treated with RU486 in adulthood compared to elav-GS/+ controls, Cox hazard analysis p<0.0001, chi^2^=89.7, n=361-377 per genotype. (D) Mortality is decreased by *elav-GS>dilp1* overexpression in the *dilp1-dilp2* double mutant background treated with RU486 in adulthood compared to elav-GS/+ controls. (E) *Dilp2>dilp1* rescue in the *dilp1-dilp2* double mutant background increases *Akh* expression compared to dilp2-Gal4/+ controls, t test p<0.001, n=5-6 per genotype. (F) Model for epistasis and interactions between *dilp1* and *dilp2* in regulating lifespan. IPCs, insulin producing cells; CC, corpora cardiaca; CA, corpora allata; JH, Juvenile Hormone.

DILP1 functions as a pro-longevity factor – it is necessary and sufficient for mutants of *dilp2* to extend longevity. This result could be readily explained if DILP1 peptide acts to inhibit the insulin receptor and its tyrosine kinase activity. DILP1 might then slow aging by decreasing insulin/IGF signaling, although our observed patterns of FOXO target gene expression are not consistent with this hypothesis. To further evaluate the potential for DILP1 to act as an IIS inhibitor, we measured body mass of adults when *dilp1* was induced as a transgene in the double mutant background. When driven by dilp2-GAL4, flies were the same mass as control flies, contrary to the expectation that an insulin/IGF signaling inhibitor would decrease body mass (Fig S5A, Supporting Information). In addition, when UAS-*dilp1* is driven by dilp2-GAL4, there was no decrease in Akt or ERK phosphorylation, again inconsistent with DILP1 acting as an IIS inhibitor (Fig S5B-D, Supporting Information). These data make it unlikely that DILP1 extends longevity by functioning as an insulin/IGF receptor inhibitor, and opens the possibility that DILP1 acts as an insulin signaling agonist that somehow induces unique signaling to assure longevity.

## Discussion

Based on mutational analyses of the insulin receptor (*daf-2, InR*) and its associated adaptor proteins and signaling elements, numerous studies in *C. elegans* and *Drosophila* established that decreased insulin/IGF signaling (IIS) extends lifespan (Clancy et al., 2001; Kenyon, Chang, Gensch, Rudner, & Tabtiang, 1993; Tatar et al., 2001). Studies on how reduced IIS in *Drosophila* systemically slows aging also reveal systems of feed-back where repressed IIS in peripheral tissue decreases DILP2 production in brain insulin producing cells (IPC), which may then reinforce a stable state of longevity assurance (Bai et al., 2012; Hwangbo et al., 2004; Wang et al., 2005). Here we find that expression of *dilp1* from the IPC is required for loss of *dilp2* to extend longevity (Fig 5C). This novel observation contrasts with conventional interpretations where reduced IIS is required to slow aging: *<u>elevated</u> dilp1* mRNA is associated with longevity in *dilp2* mutants, and transgene expression of *dilp1 <u>increases</u>* longevity.

*Dilp1* and *dilp2* are encoded in tandem, likely having arisen from a duplication event (Tatar et al., 2003). Perhaps as a result, some aspects of *dilp1* and *dilp2* are regulated in common: both are expressed in IPCs (Y. Liu et al., 2016; Rulifson et al., 2002), are regulated by sNPF (Lee et al., 2008) and have strongly correlated responses to dietary composition (Post & Tatar, 2016). Nonetheless, the paralogs are differentially expressed throughout development (Brogiolo et al., 2001). While *dilp2* is expressed in larvae, *dilp1* expression is elevated in the pupal stage when *dilp2* expression is minimal (Slaidina et al., 2009). In adults, *dilp1* expression decreases substantially after eclosion and *dilp2* expression increases (Slaidina et al., 2009).

Furthermore, as noted, DILP1 production is associated with adult reproductive diapause, (Y. Liu et al., 2016). IIS regulates adult reproductive diapause in *Drosophila,* where a somatic state is induced that prolongs survival during inclement seasons (Tatar & Yin, 2001). DILP1 may stimulate these diapause pro-longevity pathways, while expression in non-diapause adults is sufficient to extend survival even in optimal environments.

Our data suggest that *dilp1* extends longevity in part through induction of adipokinetic hormone (AKH), which is also increased during reproductive diapause (Kucerova et al., 2016) and is the functional homolog of mammalian glucagon (Bednarova, Kodrik, & Krishnan, 2013). Critically, manipulation to stimulate AKH secretion has been shown to increase *Drosophila* lifespan, potentially from catabolism of triacylglycerides and free fatty acids (Waterson et al., 2014). Here we also note that *dilp1* mutants were more sensitive to starvation than wildtype and *dilp2* mutants, further corroborating evidence that DILP1 may help mobilize nutrients during fasting (Y. Liu et al., 2016). Mammalian insulin and glucagon have inverse functions in regulating glucose storage and glycogen breakdown, and insulin decreases glucagon mRNA expression (Petersen, Vatner, & Shulman, 2017). We propose that DILP2 in *Drosophila* indirectly regulates AKH by repressing *dilp1* expression, where DILP1 otherwise induces AKH.

A further connection between *dilp1* and diapause involves juvenile hormone (JH). In many insects, adult reproductive diapause and its accompanied longevity are maintained by the absence of JH (Tatar & Yin, 2001). Furthermore, ablation of JH producing cells in adult *Drosophila* is sufficient to extend lifespan, and JH is greatly reduced in long-lived *Drosophila* insulin receptor mutants (Tatar et al., 2001; Yamamoto et al., 2013). In each case, exogenous treatment of long-lived flies with a JH analog (methoprene) restores survival to the level of wildtype or non-diapause controls. JH is a terpenoid hormone that interacts with a transcriptional complex consisting of Met (methoprene tolerant), Taimen and Kruppel homolog 1 (Kr-h1) (Jindra, Bellés, & Shinoda, 2015). As well, JH induces expression of *kr-h1* mRNA, which serves as a reliable proxy for functionally active JH. Here we find that *dilp2* mutants have reduced *kr-h1* mRNA, while the titer of this message is similar to that of wildtype in *dilp1-dilp2* double mutants. DILP1 appears to normally repress JH activity, as would occur in diapause when DILP1 is highly expressed. Such JH repression may contribute to longevity assurance during diapause as well as in *dilp2* mutant flies maintained in laboratory conditions.

Does DILP1 act as an insulin receptor agonist or inhibitor? As an inhibitor, DILP1 could interact with the insulin receptor to suppress IIS, potentially even in the presence of other insulin peptides. Such action could systemically regulate programs for longevity assurance that are stimulated by low IIS. Alternatively, DILP1 may act as an insulin receptor agonist, binding to the receptor and inducing autophosphorylation. In this case, DILP1 could stimulate signal transduction and cellular responses to confer longevity assurance that are distinct from responses associated with insulin peptides such as DILP2 or DILP5 (Post et al., 2018). To date, our limited observations on this question are ambiguous. Genetic epistasis analysis between *dilp1* and *dilp2* in terms of ERK phosphorylation suggests that DILP1 might act as an inhibitor: solo mutation of *dilp2* represses ERK phosphorylation (while increasing *dilp1* mRNA) but the double *dilp1-dilp2* mutant has wildtype ERK phosphorylation. Through a third potential mechanism, DILP1 may interact with circulating binding proteins such as IMPL2 or dALS to indirectly inhibit IIS output (Alic et al., 2011; Okamoto et al., 2013). We expect to resolve this question in a future study using synthetic DILP1 applied to cells in culture.

A precedent exists with work in *C. elegans* to suggest some insulin-like peptides might function as antagonists (Matsunaga et al., 2018; Pierce et al., 2001). *Ins-23* and *ins-18* positively regulate larval diapause and longevity (Matsunaga et al., 2018) while *ins-1* promotes dauer formation during development and longevity in adulthood (Pierce et al., 2001). Both studies suggested that the peptide chains’ altered amino acid lengths may contribute to their distinct functions. Similarly, DILP1 has an extension at the N-terminus of the B chain compared to the other DILP sequences (Brogiolo et al., 2001). Moreover, *C. elegans ins-7* suppresses *ins-6* to have contrasting and epistatic functions in particular neurons regulating olfactory learning, demonstrating a *daf-2* antagonistic role for *ins-7* (Chen et al., 2013).

While FOXO (dFOXO or DAF-16) is intimately associated with how reduced IIS regulates aging in *Drosophila* and *C. elegans* (Martins, Lithgow, & Link, 2016), we did not find evidence for this role in our epistatic analysis between *dilp1* and *dilp2* despite the robust epistatic signals for lifespan and AKH. Mutation of *dilp2* appeared to not impact FOXO activity, as measured by expression of target genes *InR* and *4eBP,* and interactions with *dilp1* did not modify this negative result. Notably, some precedence suggests only a limited role for *dfoxo* as the mediator of reduced IIS in aging, as *dfoxo* only partially rescues longevity benefits of *chico,* revealing that IIS extends lifespan through some FOXO-independent pathways (Yamamoto & Tatar, 2011). On the other hand, our whole animal analysis of dFOXO targets may obscure its role in IIS signaling if IIS and FOXO regulate aging only through actions in a few specific tissues, as previously documented (Tain et al., 2017; Wolkow, Kimura, Lee, & Ruvkun, 2000). In this vein, we find that *dilp2* controls thorax ERK signaling but not AKT, suggesting that *dilp2* mutants may activate muscle-specific ERK/MAPK anti-aging programs, although their potential action through dFOXO remains to be determined.

*Dilp1* and *dilp2* interact in a redundant fashion to regulate glycogen levels and blood sugar. These loci interact synergistically to regulate *dilp5* mRNA compensation and starvation sensitivity. In contrast, *dilp1* and *dilp2* interact in a classic epistatic fashion to modulate longevity and AKH. The distinctions and associations delineated by the classes of interactions may reflect unique ways DILP1 and DILP2 stimulate different outcomes from their common tyrosine kinase insulin-like receptor. Understanding how and what is stimulated by DILP1 in the absence of *dilp2* will likely reveal critical outputs that specific longevity assurance.

## Experimental Procedures

### Fly husbandry

Flies were reared and maintained at 25°C, 40% relative humidity and 12-hour light/dark. Adults were maintained upon agar-based diet with cornmeal (0.8%), sugar (10%), and yeast (2.5%). Fly stocks from Bloomington Stock Center include *w^1118^,dilp1* (#30880) and *w^1118^,dilp2* (#30881) mutants. *dilp2*-Gal4 stock was originally obtained from Ernst Hafen and elav-GSGal4 stock was obtained from Steven Helfand (Brown University). All stocks were backcrossed to *w^1118^* for at least five generations.

### Homologous Recombination

Homologous recombination (HR) of *dilp1* and *dilp2* in tandem was conducted as previously performed (Grönke et al., 2010; Staber, Gell, Jepson, & Reenan, 2011). See Supporting Information for detailed procedures.

### Production of UAS-*dilp1*

See Supporting Information for detailed cloning procedures. Embryos were injected with UAS-dilp1 by BestGene Inc. (thebestgene.com) yielding 5 independent transformants for *dilp1.* For this study we selected one transformant for *dilp1* that produced the strongest DILP1 immunolabeling when testing various Gal4 lines in larval and adult flies (see Fig S3).

### Lifespan assays

Two to three-day-old female adult flies, reared in density-controlled bottles and mated after eclosion, were collected with light CO_2_ anesthesia and pooled in 1 L demography cages at a density of 100-125 flies per cage. Three independent cages were used per genotype. Food vials were changed every day for the first three weeks, then every two days for the remainder of each experiment. Dead flies were removed and recorded every other day. Cox Proportional Hazard analysis was conducted in R using the “surv” package and “survdiff” function.

### RNA purification and quantitative RT-PCR

Total RNA was extracted from 20 whole mated female flies (8-10 days old) in Trizol (Invitrogen, Grand Island, NY, USA) and treated with Turbo DNase (Invitrogen). RNA was quantified with a NanoDrop ND-1000 (Thermo Fisher Scientific Inc., Wilmington, DE, USA) and reverse-transcribed with iScript cDNA synthesis (Bio-Rad Laboratories, Inc., Hercules, CA, USA). Quantitative RT-PCR was conducted with SYBR Green PCR master mix (Applied Biosystems, Carlsbad, CA, USA) and measured on an ABI prism 7300 Sequence Detection System (Applied Biosystems). mRNA abundance was calculated by comparative CT relative to ribosomal protein 49 (RP49). Primer sequences are listed in Supplemental Table S1.

### Glycogen quantification

Mated female flies aged 8-10 days were flash frozen and homogenized in PBS. Samples were heat-treated at 70°C for 10 min and spun down at 14K rpm for 3 min at 4°C. Diluted amyloglucosidase enzyme (Sigma, #A1602) was added to the glycogen standard dilutions and half of the sample wells in a 96-well clear plate. PBS was added to the glucose standards and the other half of the sample wells. After 1 hour at 37°C, Glucose Hexokinase Reagent (Thermo Scientific) was added and the plate was incubated at room temperature for 15 min. The absorbance was read at 340nm on a SpectraMax M5 platereader using Softmax Pro software. The glycogen absorbance was quantified by subtracting the glucose absorbance from the total glycogen + glucose absorbance, and normalized to protein content by BCA assay.

### Hemolymph sugar quantification

Mated female flies aged 8-10 days were decapitated to collect hemolymph in a microcentrifuge tube containing mesh. Flies were centrifuged for 4.5 min at 3K rpm and 4°C, and about 1ul hemolymph was transferred to 9ul PBS. Diluted hemolymph was heat-treated at 70°C for 5 min and transferred to tubes containing 200ul of Glucose Hexokinase Reagent (Thermo Scientific) with or without Porcine Kidney Trehalase (1:1000; Sigma-Aldrich). Reactions were incubated at 37°C for 16 hours and then transferred to wells of a clear microplate. The absorbance was read at 340nm using a SpectraMax M5 platereader and Softmax Pro software.

### Body Mass

Two females and two males in each vial were allowed to lay eggs for 24-36 hours or until proper density was attained (about 60-80 eggs). Eclosed flies were mated for two days and females were sorted to separate vials. Food was changed every other day and at 8-10 days old, flies were counted, briefly anesthetized on CO_2_ and collected in a pre-weighed microcentrifuge tube. Tubes were weighed and mass per fly was calculated.

### Western Blots

Mated female flies age 8-10 days were harvested in NP40 lysis buffer (Invitrogen) supplemented with 1mM PMSF, PhosSTOP phosphatase inhibitor cocktail (Roche #04906837001) and Protease Inhibitor Cocktail (Invitrogen). Samples were run on SDS-PAGE (Invitrogen NuPAGE). Gels were transferred to nitrocellulose membrane (Whatman) for one hour at 30V and washed in TBS-T for 10 min. Membranes were blocked with 5% BSA in TBS-T for one hour, and incubated with antibody 1:1000 in 5% BSA overnight at 4°C. Antibodies used were from Cell Signaling Technology: *Drosophila* phospho-Akt Ser505 (#4054S), Pan-Akt (#4691S), Pan-phospho-ERK (#4370S), Pan-ERK (#9102S). Blots were washed 3 times for 5 min and incubated in HRP-conjugated anti-rabbit secondary antibody (Jackson Immunoresearch) 1:5000 in 1% BSA for one hour at room temperature. Blots were washed and incubated with ECL reagent (Perkin Elmer #NEL121001EA) for 5 minutes. Blots were imaged and analyzed by volume densitometry in ImageLab (BioRad).

### Antisera and immunocytochemistry

Tissues from larvae or 7-day-old female adults were dissected in 0.1 M PBS, then fixed for four hours in ice-cold 4% paraformaldehyde (PFA), and rinsed in PBS three times for one hour. Incubation with primary antiserum was performed for 48 hours at 4°C. After rinsing in PBS with 0.25% Triton-X 100 (PBS-Tx) four times, tissues were incubated with secondary antibody for 48 hours at 4°C. After washing in PBS-Tx, tissues were mounted in 80% glycerol with 0.1 M PBS. Primary antisera used were: Rabbit antisera to DILP1 C-peptide (Y. Liu et al., 2016) at a dilution of 1:10,000, rabbit antisera to DILP2 and DILP3 A-chains (Veenstra, Agricola, & Sellami, 2008) at a dilution of 1:2000, rabbit antisera to AKH was kindly donated by M. Brown (Athens, GA) used at 1:1000, rabbit anti-GFP at 1:000 (Invitrogen, Carlsbad, CA). Secondary antisera used were: goat anti-rabbit Alexa 546 antiserum and goat anti-rabbit Alexa 488 antiserum (Invitrogen, Carlsbad, CA). at 1:1000.

### Image analysis

Confocal images were captured with a Zeiss LSM 780 confocal microscope (Jena, Germany) using a 40× oil immersion objective. The projection of z-stacks was processed using Fiji (https://imagej.nih.gov/ij/). The cell body outlines were extracted manually and the staining intensity was determined using Fiji. The background intensity for all samples was recorded by randomly selecting three small regions near the cell body of interest. The final intensity value of the cell bodies was determined by subtracting the background intensity.

### Starvation resistance assays

Flies were mated for two days and females were separated and maintained for 6-7 days on normal food. For fasting, the flies were kept in vials containing 5 ml of 0.5% agarose (Sigma-Aldrich). Dead flies were counted at least every 12 hours. No less than 118 flies from 3 replicates were used for the analysis.

## Acknowledgements

We thank Barry Pfeiffer for sharing the vector used to initiate production of the UAS-*dilp1* transgenic fly. Funding for SP, RY and MT was supported by NIH R37 AG024360. SP was additionally supported by NIH T32 AG 41688-3 and by AFAR GR5290420. SL and DRN were supported by The Swedish Research Council (Vetenskapsrådet 2015-04626). JAV was supported by institutional funds from the CNRS.

## Author Contributions

SP and MT designed experiments and interpreted results. SP executed experiments. SF and RY executed experiments and contributed to experimental design. DN contributed to experimental design and interpretation of results. JV contributed to developing reagents. SP and MT wrote the manuscript and all authors edited the manuscript.

## Supporting Information

**Table S1. Primer Sequences**

**Fig S1. Validation of *dilp1-dilp2* mutants.**

**Fig S2. Additional epistasis analyses between *dilp1* and *dilp2* single and double mutants.**

**Fig S3. UAS-*dilp1* expression drives peptide production.**

**Fig S4. UAS-*dilp1* expression does not repress insulin/IGF signaling or growth.**

**Data S1. Supplemental Methods and Figure Legends**

**Table S1. Primer Sequences**

